## Supplementary Material for "Absorption and fixation times for evolutionary processes on graphs"

The objective of this Supplementary Material is twofold, first to provide a mathematical basis for the results of the main text and second to illustrate these results with a graphical description for some classes of graphs and low-order graphs.

### S1. Absorption and fixation times for finite-state Markov chains with two absorbing states

In this section we slightly change the notation with respect to the main text, hoping that the formulas will be easier to read. We consider a finite-state discrete-time Markov chain $\{S_n\}_{n\geq 0}$ with transition matrix $P$, whose state space is divided into three communicating classes (the generalization to more classes is straightforward): $a$ and $b$ are absorbing states and $C$ is the class of all transient states, that we enumerate $i = 1, 2, \ldots, m$.

Here we provide closed matrix expressions for the (mean) times the process takes until it is absorbed by a given absorbing state, starting in a transient state $i \in C$. As it is well known, we can order the states in such a way that the transition matrix of the process can be written in the form:

$$P = \left[\begin{array}{c|c} Q & R \\ \hline 0 & I \end{array}\right],$$

where $Q$ is the $m \times m$ matrix whose entries are the transition probabilities between transient states:

$$Q = \begin{bmatrix} p_{1,1} & p_{1,2} & \cdots & p_{1,m} \\ p_{2,1} & p_{2,2} & \cdots & p_{2,m} \\ \vdots & \vdots & \ddots & \vdots \\ p_{m,1} & p_{m,2} & \cdots & p_{m,m} \end{bmatrix}$$

and $R$ is the $m \times 2$ matrix of transition probabilities from transient to absorbing states:

$$R = \begin{bmatrix} p_{1,a} & p_{1,b} \\ p_{2,a} & p_{2,b} \\ \vdots & \vdots \\ p_{m,a} & p_{m,b} \end{bmatrix}.$$

In what follows we adapt the notation and terminology to the context of evolutionary models on graphs:

- $a$ is the mutant *extinction* state,
- $b$ is the mutant *fixation* state,
- $\mathcal{P}_{i,a}$ is the set of all trajectories that start in state $i \in C$ and are absorbed by state $a$,
- $\mathcal{P}_{i,b}$ is the set of all trajectories that start in state $i \in C$ and are absorbed by state $b$,

- $\Phi_{i,a} = \mathbb{P}[\mathcal{P}_{i,a}]$ is the probability of extinction given that the process started in state $i \in C$,
- $\Phi_{i,b} = \mathbb{P}[\mathcal{P}_{i,b}]$ is the probability of fixation given that the process started in state $i \in C$,
- $T_{i,a}$ is the extinction time given that the process started in state $i \in C$,
- $T_{i,b}$ is the fixation time given that the process started in state $i \in C$,
- $\tau_i$ is the absorption time given that the process started in state $i \in C$,

With these quantities, which we will shortly explain how they are calculated, we build the following matrices:

$$\Phi = \begin{bmatrix} \Phi_{1,a} & \Phi_{1,b} \\ \Phi_{2,a} & \Phi_{2,b} \\ \vdots & \vdots \\ \Phi_{m,a} & \Phi_{m,b} \end{bmatrix} \quad , \quad T = \begin{bmatrix} T_{1,a} & T_{1,b} \\ T_{2,a} & T_{2,b} \\ \vdots & \vdots \\ T_{m,a} & T_{m,b} \end{bmatrix} \quad , \quad \tau = \begin{bmatrix} \tau_1 \\ \tau_2 \\ \vdots \\ \tau_m \end{bmatrix}$$

**Computation of $\Phi$.** As it is well known, we have:

$$\Phi = (I - Q)^{-1} R, \tag{S1.1}$$

where $I$ is the identity matrix $m \times m$. The matrix

$$(I - Q)^{-1} = \sum_{n=0}^{\infty} Q^n$$

is called the *fundamental matrix.* For $i \in C$ and $\alpha = a$ or $b$, the entry $[Q^n R]_{i,\alpha}$ is the probability of all the trajectories that remain for $n$ transitions in class $C$ starting in state $i$ and ending in absorbing state $\alpha$.

**Computation of $T$.** The absorption time, starting the process in state $i \in C$, is the random variable

$$\tau_i = \min\{\, n \geq 1 \mid X_n = a \text{ or } X_n = b \,\}.$$

As it is also well known, the expected absorption time vector $\mathbb{E}[\tau]$ with entries $\mathbb{E}[\tau_i]$ is given by

$$\mathbb{E}[\tau] = (I - Q)^{-1} \mathbb{1}, \tag{S1.2}$$

where $\mathbb{1}$ is the vector with all entries equal to 1.

For computing the expected fixation (resp. extinction) time we must condition the process to fixation (resp. extinction):

$$\begin{aligned} T_{i,\alpha} &= \mathbb{E}\left[\tau_i | \mathcal{P}_{i,\alpha}\right] \\ &= \sum_{n=1}^{\infty} n\, \mathbb{P}\left[\tau_i = n | \mathcal{P}_{i,\alpha}\right] \\ &= \sum_{n=1}^{\infty} n\, \frac{\mathbb{P}\left[\tau_i = n, X_{\tau_i} = \alpha\right]}{\mathbb{P}\left[\mathcal{P}_{i,\alpha}\right]} \\ &= \frac{\sum_{n=1}^{\infty} n\, [Q^{n-1} R]_{i,\alpha}}{\Phi_{i,\alpha}}. \end{aligned}$$

Recall that the entry $[Q^{n-1}R]_{i,\alpha}$ means that the process remains $n-1$ transitions in class $C$ and ends in the absorbing state $\alpha$.

So, we can write:

$$T_{i,\alpha} = \frac{\left[\left(\sum_{n=1}^{\infty} n\, Q^{n-1}\right) R\right]_{i,\alpha}}{\left[(I - Q)^{-1} R\right]_{i,\alpha}}.$$

Finally we note:

$$(I-Q)^{-2} = \sum_{l=0}^{\infty}\sum_{m=0}^{\infty} Q^{l+m} = \sum_{n=1}^{\infty} n\, Q^{n-1},$$

and then

$$T_{i,\alpha} = \frac{\big[(I-Q)^{-2}R\big]_{i,\alpha}}{[(I-Q)^{-1}R]_{i,\alpha}} \tag{S1.3}$$

that means we can obtain all the expected fixation and extinction times from the fundamental matrix. This equation has been used for the exact computation of the mean fixation times on low order graphs (see Section S4). Finally, we can observe:

$$\mathbb{E}[\tau_i] = \Phi_{i,a}\, T_{i,a} + \Phi_{i,b}\, T_{i,b} \tag{S1.4}$$

for all $i \in C$.

### S2. Embedded jump process and Doob's transform

Let P be any of processes considered here, namely Moran process and Bernoulli or binomial proliferation processes, defined on a connected undirected graph $G = (V, E)$. In all cases, the state space is the power set $\mathcal{S} = \mathcal{P}(V)$ formed of all subsets of $V$ and the process is defined by a $(m+2) \times (m+2)$ matrix $P$ whose entries are the transition probabilities $p_{S,S'}$ between two states $S$ and $S'$ and $m+2 = 2^N$. Without loss of generality, adding all inessential classes (in the sense of [9]) to $C$, we can assume that there are tree communicating classes as supposed in Section S1: $\emptyset$ and $V$ are absorbing states and $C = \mathcal{S} - \{\emptyset, V\}$ is the class of all transient states. In fact, as described in Section S1, we could indeed consider any finite-state discrete-time Markov chain $\mathrm{P} = \{S_n\}_{n\geq 0}$ with similar properties. Note however that states belong to different communicating classes have an important role in the study of $G$-symmetry introduced in the main text.

As usual, the process P can be identified to the random walk on the state space $\mathcal{S}$ endowed with the natural graph structure derived from the transition matrix $P$. Recall that two states $S$ and $S'$ are connected by an oriented edge if and only if $p_{S,S'} > 0$ and this edge is naturally labelled by the transition probability. In other words, the space of trajectories $\mathcal{P} = \{\sigma = \{S_n\}_{n\geq 0} : \sigma(n) = S_n \in \mathcal{S}\}$ is endowed with the probability measure

$$\mathbb{P}[\sigma(0) = S_0, \ldots, \sigma(n) = S_n] = \delta_{S_0}(\sigma(0)) p_{S_0,S_1} \cdots p_{S_{n-1},S_n}$$

where $S_0, \ldots, S_n \in \mathcal{S}$ and the Dirac measure $\delta_{S_0}$ is the initial distribution of $\mathbb{P}$. We shall denote by $\mathbb{P}_{S_0}$ the probability conditioned to the initial state $S_0$.

**Embedded jump process.** The *embedded jump process* $\hat{\mathrm{P}}$ on $G$ is given by defining

$$\hat{p}_{S,S'} = \frac{p_{S,S'}}{1 - p_{S,S}} \tag{S2.1}$$

for any pair of non-absorbing states $S, S' \in \mathcal{S} - \{\emptyset, V\}$. The state space $\hat{\mathcal{S}}$ remains equal to $\mathcal{S}$, but the graph structure of $\hat{\mathcal{S}}$ is obtained from that of $\mathcal{S}$ by removing loops in all transient states. The trajectory space $\mathcal{P}$ retracts onto $\hat{\mathcal{P}}$ by sending each trajectory $\sigma \in \mathcal{P}$ to the trajectory $\hat{\sigma} \in \hat{\mathcal{P}}$ with non-trivial transitions, that is, such that $\hat{\sigma}(n) \neq \hat{\sigma}(n+1)$ for all $n \geq 0$. This retraction projects the measure $\mathbb{P}$ to the measure $\hat{\mathbb{P}}$ given by

$$\begin{aligned}\hat{\mathbb{P}}[\hat{\sigma}(0) = S_0, \ldots, \hat{\sigma}(n) = S_n] &= \delta_{S_0}(\hat{\sigma}(0)) \hat{p}_{S_0,S_1} \cdots \hat{p}_{S_{n-1},S_n} \\ &= \delta_{S_0}(\hat{\sigma}(0)) \frac{p_{S_0,S_1}}{1 - p_{S_0,S_0}} \cdots \frac{p_{S_{n-1},S_n}}{1 - p_{S_{n-1},S_{n-1}}}\end{aligned} \tag{S2.2}$$

From Equation S2.2 and conditioning to the initial state $S_0$, we deduce that the fixation probabilities of $S_0$ coincide for both processes P and $\hat{\mathrm{P}}$. Indeed, if we denote $\mathcal{F}_{S_0}$ and $\hat{\mathcal{F}}_{S_0}$ the fixation events, both conditioned to the initial state, then

$$\hat{\Phi}_{S_0} = \hat{\mathbb{P}}_{S_0}\left[\exists n \geq 0 : \hat{S}_n = V \mid \hat{S}_0 = S_0\right] = \hat{\mathbb{P}}_{S_0}(\hat{\mathcal{F}}_{S_0})$$

and

$$\Phi_{S_0} = \mathbb{P}_{S_0}\left[\exists n \geq 0 : S_n = V \mid S_0 = S_0\right] = \mathbb{P}_{S_0}(\mathcal{F}_{S_0})$$

are equal.

The average absorption times $\tau_{S_0}$ and $\hat{\tau}_{S_0}$ (which are defined on the trajectory space $\mathcal{P}_{S_0}$ and $\hat{\mathcal{P}}_{S_0}$ conditioned to same initial state $S_0$) are related by

$$\begin{aligned}
\tau_{S_0}(\sigma) &= n + \sum_{i=0}^{n-1} \frac{p_{S_i,S_i}}{1 - p_{S_i,S_i}} \\
&= n + \sum_{i=0}^{n-1} \big(\frac{1}{1 - p_{S_i,S_i}} - 1\big) \qquad \text{(S2.3)} \\
&= \sum_{i=0}^{n-1} \frac{1}{1 - p_{S_i,S_i}}
\end{aligned}$$

where $n = \hat{\tau}_{S_0}(\hat{\sigma})$ is the number of non-trivial transitions from $\sigma(0) = S_0$ to $\sigma(n) = S_n$.

The expected fixation time

$$T_{S_0} = \mathbb{E}[\tau_{S_0} | \mathcal{F}_{S_0}]$$

is bounded by the product of the expected number of non-trivial transitions and the maximum mean sojourn time in transient states:

$$T_{S_0} \leq \mathbb{E}[\hat{\tau}_{S_0} | \hat{\mathcal{F}}_{S_0}]\, max\big\{\frac{1}{1 - p_{S,S}} : S \in C\big\}.$$

**Remark S1.** The proof of the Circulation Theorem in [5] implicitly used the embedded jump process by associating to any *circulation* on $G$ (that is, assuming the temperature of any vertex is constant, equal to 1) the random walk on the integer interval $[0, N]$ with absorbing states 0 and $N$ and forward bias $r$. Indeed, for any state $S$, the probability of increasing or decreasing the number of mutant is equal to

$$\frac{rw_+(S)}{rw_+(S) + w_-(S)} \quad \text{or} \quad \frac{w_-(S)}{rw_+(S) + w_-(S)}$$

where $w_+(S)$ and $w_-(S)$ are the leaving and entering weights of $S$. For a circulation on $G$, both weights are equal, depending only on $|S| = i$. Compare to the embedded jump process on the complete graph described in Figure 1 below. A method to compute $\hat{\tau}_{S_0}$ and $\hat{T}_{S_0}$ using Wald's martingales has been applied to isothermal graphs in [8].

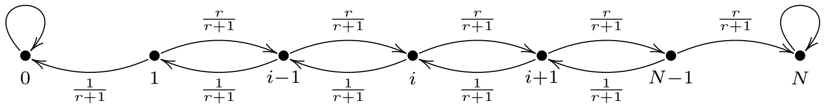


FIGURE 1. The embedded jump process on the complete graph $K_N$.

**Doob transform.** Given any absorbing state of the process P, we can replace the conditioned process by the *Doob transform* $\check{\mathrm{P}}$ as explained in [4, Chapter 17]. For simplicity, we will restrict to the fixation event.

For any pair of non-absorbing states $S, S' \in \mathcal{S}$, we define the transition probability

$$\check{p}_{S,S'} = \frac{\Phi_{S'}}{\Phi_S} p_{S,S'}, \tag{S2.4}$$

where $\Phi_S$ and $\Phi_{S'}$ are the fixation probabilities of states $S$ and $S'$. The graph structure of the state space $\check{\mathcal{S}}$ is the same that of the original process $\mathcal{S}$, but the transition probabilities are biased to fixation. Observe that all sojourn times remain unchanged since

$$\check{p}_{S,S} = \frac{\Phi_S}{\Phi_S} p_{S,S} = p_{S,S}.$$

By construction, the expected fixation times are also equal:

$$T_{S_0} = \mathbb{E}[\tau_{S_0} | \mathcal{F}_{S_0}] = \mathbb{E}[\check{\tau}_{S_0}]. \tag{S2.5}$$

This equality can be also derived from the following identity:

$$\begin{aligned} \check{\mathbb{P}}_{S_0}[\check{\tau}_{S_0} = n] &= \sum_{S_1,\ldots,S_{n-1}} \check{p}_{S_0,S_1} \cdots \check{p}_{S_{n-1},S_n} \\ &= \sum_{S_1,\ldots,S_{n-1}} \frac{\Phi_{S_1}}{\Phi_{S_0}} p_{S_0,S_1} \cdots \frac{\Phi_{S_n}}{\Phi_{S_{n-1}}} p_{S_{n-1},S_n} \qquad \text{(S2.6)} \\ &= \frac{1}{\Phi_{S_0}} \mathbb{P}_{S_0}[\tau_{S_0} = n] \end{aligned}$$

as $S_n = V$ and $\Phi_{S_n} = 1$. Indeed, now we have:

$$\begin{aligned} \mathbb{E}[\check{\tau}_{S_0}] &= \sum_{n=1}^{\infty} n \check{\mathbb{P}}_{S_0}[\check{\tau}_{S_0} = n] \\ &= \frac{1}{\Phi_{S_0}} \sum_{n=1}^{\infty} n \mathbb{P}_{S_0}[\tau_{S_0} = n] \\ &= \frac{1}{\Phi_{S_0}} \mathbb{E}[\tau_{S_0}] = \mathbb{E}[\tau_{S_0} | \mathcal{F}_{S_0}]. \qquad \text{(S2.7)} \end{aligned}$$

**Proposition S1.** *The embedded jump construction and the Doob transform commute.*

*Proof.* Indeed, for any pair of non-absorbing states $S$ and $S'$, we wave:

$$\begin{aligned} \hat{\check{p}}_{S,S'} &= \frac{\check{p}_{S,S'}}{1 - \check{p}_{S,S}} \\ &= \frac{\frac{\Phi_{S'}}{\Phi_S} p_{S,S'}}{1 - p_{S,S}} \\ &= \frac{\Phi_{S'}}{\Phi_S} \frac{p_{S,S'}}{1 - p_{S,S}} \\ &= \frac{\hat{\Phi}_{S'}}{\hat{\Phi}_S} \hat{p}_{S,S'} = \check{\hat{p}}_{S,S'} \end{aligned}$$

as we wanted. □

Now we can prove Proposition 2.1 of the main text (which can also be derived directly from the matrix identities (S1.2) and (S1.3)):

*Proof of Proposition 2.1.* As we already observed the fixation probabilities $\Phi_{S_0}$ and $\hat{\Phi}_{S_0}$ of the processes P and $\hat{\mathrm{P}}$ are equal and then their limit are also equal:

$$\lim_{r\to+\infty} \Phi_{S_0} = \lim_{r\to+\infty} \hat{\Phi}_{S_0}.$$

Both process depend only on $r \in (0, +\infty)$ if $\mathrm{P} = \mathrm{P}^M$ is the Moran process, and also on $p \in (0, 1]$ if $\mathrm{P} = \mathrm{P}^\beta$ is a proliferation process. For the complete graph $G = K_N$, it is evident that the process $\hat{\mathrm{P}}$ approaches the *forward-only process* (described in Figure 2(c)) when $r \to +\infty$. In general, for any graph $G$, the limit process does not have trajectories that lead to extinction. It follows that

$$\lim_{r\to+\infty} \Phi_{S_0} = \lim_{r\to+\infty} \hat{\Phi}_{S_0} = 1.$$

On the other hand, combining Equations (S2.5) and (S2.7), we deduce:

$$\begin{aligned}
\lim_{r\to+\infty} T_{S_0} &= \lim_{r\to+\infty} \mathbb{E}[\check{\tau}_{S_0}] \\
&= \lim_{r\to+\infty} \frac{1}{\Phi_{S_0}} \mathbb{E}[\tau_{S_0}] \\
&= \lim_{r\to+\infty} \mathbb{E}[\tau_{S_0}] \\
&= \lim_{r\to+\infty} \tau_{S_0}.
\end{aligned}$$

We use here the same abusive notation $\tau_{S_0}$ for the expected absorption time in the main text, having the same dependence on $r$ and $p$ as before. □

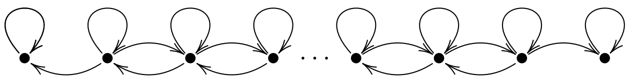

(a) Moran process

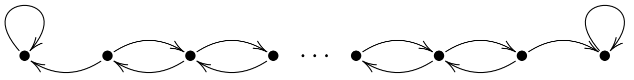

(b) Embedded jump process

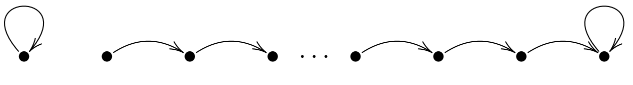

(c) Forward-only process

Figure 2. Moran, embedded jump and forward-only processes on a complete graph.

### S3. Graphical model and red process

As in the previous section, let P be any process $\mathrm{P}^M$ or $\mathrm{P}^\beta$ with $\beta = B, b$ on a connected undirected graph $G = (V, E)$ with fitness $r \in (0, +\infty)$ and proliferation parameter $p \in (0, 1]$. According to [1], such a process can be realised from a graphical model similar to the Harris' representation for contact processes [2]. This realisation is done on the set $G \times [0, +\infty)$ by taking two independent Poisson distributions

$$\Pi_\varepsilon(v) = \{\tau_i^\varepsilon(v) : i \in \mathbb{N}^*\}$$

on $[0, +\infty)$, with rate $r$ for $\varepsilon = 1$ and rate 1 for $\varepsilon = 2$. Each distribution determines a set of coloured marks

$$\mathcal{M}_\varepsilon = \{(v,\tau) \in V \times [0,+\infty) : \tau \in \Pi_\varepsilon(v)\},$$

namely red for $\varepsilon = 1$ and blue for $\varepsilon = 2$. Both sets are completed by selecting the corresponding sets of red and blue arrows. For any $(v,\tau) \in V \times [0,+\infty)$, we denote $A(v,\tau) \subset E \times \{v\}$ the set of edges with origin $v$. The set $\mathcal{A}_1$ of red arrows is defined as follows:

1) For the Moran process, an element of $A(v,\tau)$ will be chosen at random if $(v,\tau) \in \mathcal{M}_1$.
2) For Bernoulli proliferation, we will define $\mathcal{A}_1(v,\tau) = A(v,\tau)$ with probability $p$ and $\mathcal{A}_1(v,\tau) = \emptyset$ with probability $1-p$.
3) For binomial proliferation, we will choose randomly and independently each element of $\mathcal{A}_1(v,\tau)$ within $A(v,\tau)$ with probability $p$.

To define the set $\mathcal{A}_2$ of blue arrows, we will proceed similarly to that happens in the Moran process by choosing at random an unique element of $A(v,\tau)$ if $(v,\tau) \in \mathcal{M}_2$.

This graphical model provides a continuous-time Markov chain $\{X_t\}_{t\in[0,+\infty)}$ realising the discrete-time Markov chain $\{S_n\}_{n\in\mathbb{N}}$ defined by P. We will use the same notation for the expected absorption and fixation times of both processes, even though they differ by a factor that depends on fitness. Ignoring blue marks and arrows, we obtain another continuous-time Markov chain $\{\bar{X}_t\}$, which we will call the *red process* associated to $\{X_t\}$. In fact, up to scale, this process is uniquely determined by the underlying graph.

Processes $\{X_t\}$ and $\{X'_t\}$ having the same fitness $r' = r$, but corresponding to different proliferation parameters $0 < p' < p$ can be coupled taking the same random set of red and blue marks, but randomly deleting red arrows. Indeed, repeating the procedure described in [1], the sets $\mathcal{M}'_1$ and $\mathcal{M}'_2$ of red and blue marks are equal to $\mathcal{M}_1$ and $\mathcal{M}_2$. For the blue arrows, we have also $\mathcal{A}'_2 = \mathcal{A}_2$, but $\mathcal{A}'_1$ is obtained from $\mathcal{A}_1$ by deleting red arrows with probability $1 - p'/p$ by using Bernoulli or binomial draws. If the overall processes $\{X_t\}$ and $\{X'_t\}$ are coupled, then the red processes $\{\bar{X}_t\}$ and $\{\bar{X}'_t\}$ are also coupled.

Choosing a vertex $v \in V$ and conditioning to the state $X_0 = S_0 = \{v\}$, we define the first sojourn time as

$$t_1 = \inf\{t \in [0,+\infty) : X_t \neq X_0\}$$

Recursively, assuming $t_n < +\infty$, we define

$$t_{n+1} = \inf\{t \in [t_n,+\infty) : X_t \neq X_{t_n}\}$$

The distribution of $t_1$ conditioned to $X_0 = S_0 = \{v\}$ is exponential with parameter $\lambda(S_0)$. In the same way the distribution of the sojourn time

$$s_n = t_n - t_{n-1}$$

conditioned to $X_{t_{n-1}} = S_{n-1}$ is exponential with parameter $\lambda(S_{n-1})$. Therefore, if $\{S_n\}$ is the embedded jump chain associated to $\{X_t\}$, the distribution of the sum

$$t_n = s_1 + \cdots + s_n$$

is a hypoexponential distribution with parameters $\lambda(S_0), \ldots, \lambda(S_{n-1})$ when we conditioned to $X_0 = S_0, \ldots, X_{t_{n-1}} = S_{n-1}$. A related description can be seen in [3, Section 4.8].

Associated to the random variable

$$\mathbf{n} = \sup\{n \geq 1 : t_n < +\infty\},$$

we introduce a new random variable

$$t_{\mathbf{n}} = \sum_{n=1}^{\infty} t_n \mathbb{1}_{\{\mathbf{n}=n\}} = \lim_{m\to+\infty} \sum_{n=1}^{m} t_n \mathbb{1}_{\{\mathbf{n}=n\}} \tag{S3.1}$$

where $\mathbb{1}_{\{\mathbf{n}=n\}}$ is the characteristic function of the event $\{\mathbf{n} = n\}$. By definition, the expected absorption time $\tau_{S_0} = \mathbb{E}[t_{\mathbf{n}}]$. The same construction applies to the red process $\{\bar{X}_t\}$ conditioning to the state $\bar{X}_0 = S_0 = \{v\}$, and all notations will be consequently modified by adding a bar.

For the red process, assuming all random variables $\bar{s}_n$ are identically distributed, we deduce from Wald's first lemma that, for any state $S_0 = \{v\}$, the expected fixation time

$$\bar{T}_{S_0} = \mathbb{E}[\bar{t}_{\bar{\mathbf{n}}}] = \mathbb{E}[\bar{t}_1]\,\mathbb{E}[\bar{\mathbf{n}}]. \tag{S3.2}$$

Finally, observe that the expected fixation time $\bar{T}_{S_0}$ of the (neutral) red process is equal to the asymptotic value $\lim_{r\to+\infty} T_{S_0}$ of the expected fixation time $T_{S_0}$ of the overall process when the fitness $r > 0$ goes to infinity. In particular, this limit always exists.

**Remark S2.** The computation for complete, cycle and star graphs included in the main text suggests that $\lim_{r\to+\infty} T_{S_0}$ is related with the *cover time* of the simple random walk (SRW in short) on the underlying graph, see [4] and [6]. For both times, the initial state $S_0$ can be replaced by the uniform distribution, the stationary distribution, or the worst-case initial state. In fact, the (neutral) red process can be interpreted as a specific cover process to be compared to the SRW. We know that both processes have the same cover time for complete and cycle graphs, but SRW is more efficient in covering star graphs.

In the overall process, assuming again that the random variable $s_n$ are identically distributed and conditioning to the fixation event $\mathcal{F}_{S_0}$, we can reformulate (S3.2) as follows:

$$T_{S_0} = \mathbb{E}[t_{\mathbf{n}}|\mathcal{F}_{S_0}] = \mathbb{E}[t_1|\mathcal{F}_{S_0}]\,\mathbb{E}[\mathbf{n}|\mathcal{F}_{S_0}]. \tag{S3.3}$$

In fact, the expected absorption time (conditioned to $X_0 = S_0 = \{v\}$) admits a related description from the sequence of expected harmonic mean $H_n$ of parameters $\lambda(S_0), \dots, \lambda(S_{n-1})$. Indeed, as

$$\mathbb{E}[t_n] = \frac{n}{H_n},$$

we deduce

$$\begin{aligned} \mathbb{E}[t_{\mathbf{n}}] &= lim_{m\to+\infty} \sum_{n=1}^{m} \mathbb{E}[t_n \mathbb{1}_{\{\bar{\mathbf{n}}=n\}}] \\ &= lim_{m\to+\infty} \sum_{n=1}^{m} \frac{1}{H_n} \mathbb{E}[n \mathbb{1}_{\{\mathbf{n}=n\}}] \\ &= lim_{m\to+\infty} \sum_{n=1}^{m} A_n \mathbb{E}[\mathbf{n} \mathbb{1}_{\{\mathbf{n}=n\}}] \end{aligned} \tag{S3.4}$$

where $A_n$ is the expected arithmetic mean of $\frac{1}{\lambda(S_0)}, \dots, \frac{1}{\lambda(S_{n-1})}$.

However, the conditions imposed to the sojourn times to obtain (S3.2) and (S3.3) are too restrictive, and (S3.4) is too imprecise to obtain any kind of monotonicity. In the main text, we use a recursive argument to prove the monotonicity in $p$ of the expected sojourn times for the red process associated to proliferation processes on graphs.

### S4. Exact computation in low order

Initially all analytic computations for complete, cycle and star graphs suggested that even if the proliferation is not advantageous for the fixation of mutant individuals –leaving unchanged the probability of take over the whole population–, once that this happen, the expected time for the fixation is always reduced in both Bernoulli and binomial proliferation processes. See Section 3 and 4 of the main text.

To test this idea, using the same method to that described in [1], we have developed a Python package, written in C++, that computes the values of expected absorption and fixation times with high precision for all graphs of order 6. From equations (S1.2) and (S1.3), we can easily deduce that $\tau_1^\beta$ and $T_1^\beta$ (for $\beta = b, B$) are rational functions on the fitness $r$ and the proliferation parameter $p$. The degree of both denominator and numerator polynomials is bounded by $2^N - 2$ (see Section S1). But as we have already seen, this limit can be reduced using the symmetries of the process, that is, identifying equivalent states under automorphisms of the process.

Thus, given any graph $G$ and any fitness value $r \in \mathbb{Q}$, the main steps are as follows:

1) We compute generators for the graph automorphism group.
2) We identify equivalent states with graph automorphisms and Breadth-First Search for accessible states. We reduce the size of the transition matrices.
3) We compute the transition matrices with coefficients in $\mathbb{Q}[r, p]$ for both the standard Moran process and the one with proliferation $\beta$ implemented. Their size is limited by the number of accessible states, improving the computation time.
4) We compute the fixation probability $\Phi_1^M(r)$, the absorption time $\tau_1^M(r)$ and the fixation time $T_1^M(r)$ for Moran process with Gaussian elimination.
5) To compute the coefficients $a_i$ and $b_i$ of denominator and numerator of the rational function $\Phi_1(r, p_c^\beta) = \Phi_1^M(r)$ it is enough to compute $\Phi_1^\beta(r, p_i)$ for enough values $p_i = 1/i$. Thus we get a linear system on $a_i$ and $b_i$ that can be solved by Gaussian elimination. This system is usually underdetermined since the degree may be overestimated and hence there are multiple writings for the same rational function. Then we reduce the degree accordingly and recompute this step.
6) We apply bisection algorithm to find $p_c^\beta$ in the interval $[0, 1]$. We make up to 53 divisions to get an error near machine epsilon. The rational value obtained above is converted to a floating point double and return.
7) Finally, the times $\tau_1(r, p_c^\beta)$ and $T_1(r, p_c^\beta)$ are computed solving the corresponding linear system with rational coefficients.

This algorithm allows us to compute, with high precision, the absorption and fixation times for all graphs of order 6 for both type of proliferation updating.

In Figures 3(a) and 3(c) the mean absorption time $\tau_1^\beta(r, p_c)$ is shown for fitness values $r$ between 0.1 to 10 and critical proliferation parameter $p_c$ for all graphs of order 6. Complete, cycle and star graphs are specifically marked, as well as trees and isothermal graphs. Similarly the mean fixation time $T_1^\beta(r, p_c)$ is shown in Figures 3(b) and 3(d). As we have proved in the main text, unless for the complete and cycle graph, the G-symmetry shown in Figures 3(b) and 3(d) for the expected fixation time under Moran updating is only apparent. A detailed description of the fitness values for which mean absorption and fixation times are maximum, as well the values of these maximum times, can be found in the site [7].

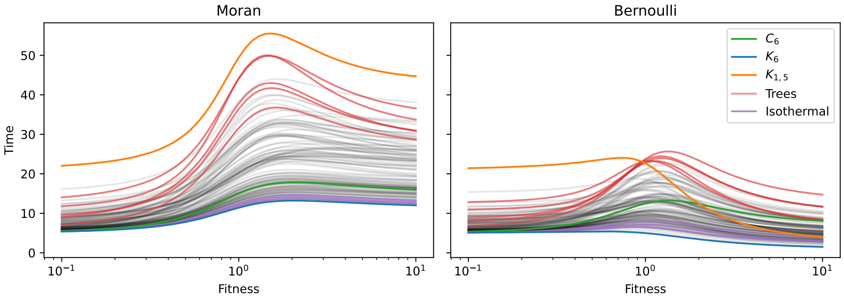


(a) Absorption times $\tau_1^M(r)$ and $\tau_1^B(r, p_c)$

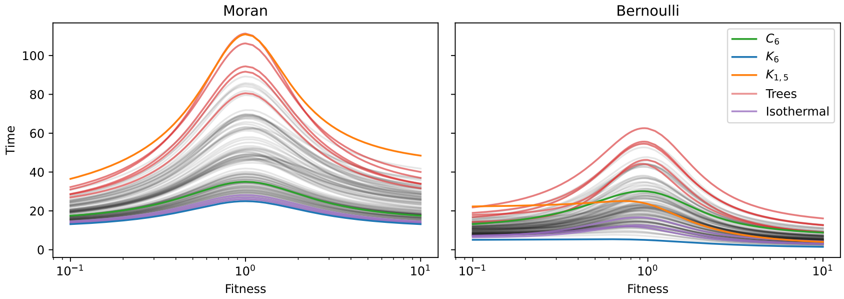


(b) Fixation times $T_1^M(r)$ and $T_1^B(r, p_c)$

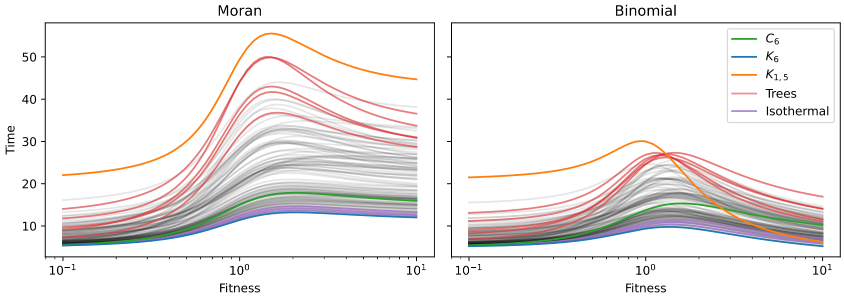


(c) Absorption times $\tau_1^M(r)$ and $\tau_1^b(r, p_c)$

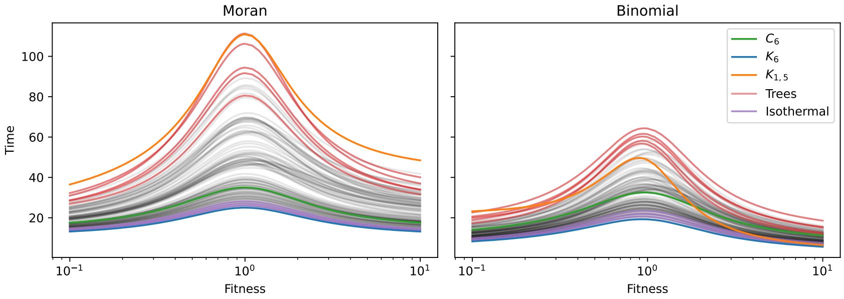


(d) Fixation times $T_1^M(r)$ and $T_1^b(r, p_c)$

FIGURE 3. Comparing mean times for non-proliferating (Moran process) and proliferating (Bernoulli and binomial) mutant individuals: (a)-(c) absorption times and (b) -(d) fixation times for all graphs of order 6.

In Figure 4, we show the rates

$$\rho^\beta(r,p) = \frac{\tau_1^M(r)}{\tau_1^\beta(r,p)} \quad \text{and} \quad P^\beta(r,p) = \frac{T_1^M(r)}{T_1^\beta(r,p)}$$

for all graphs of order 6. Fitness values again range between 0.1 to 10. Thus, at least in low order, we can graphically visualize the asymptotic properties of these rates which have been exhibited in Sections 3 and 4 of the main text. It should be noted that cycle and line graphs have the lowest rate, while the star graph has the greater rate from a certain fitness value. The asymmetry in the Moran process for certain graphs seems also to appear in proliferation processes, at least in the critical regime $p = p_c$.

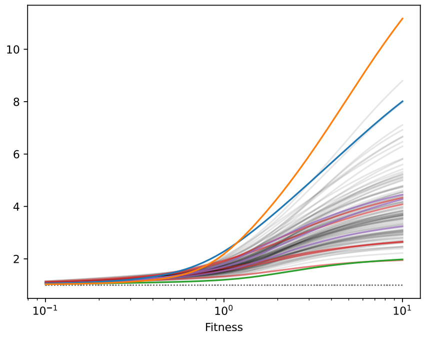


(a) Absorption time rate $\rho^B(r,p_c)$

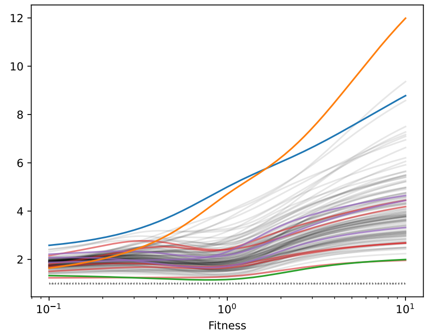


(b) Fixation time rate $P^B(r,p_c)$

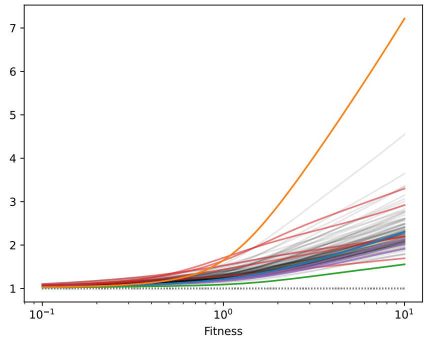


(c) Absorption time rate $\rho^b(r,p_c)$

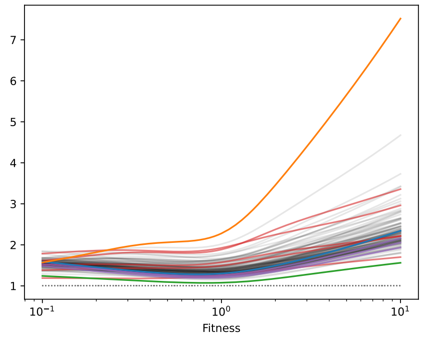


(d) Fixation time rate $P^b(r,p_c)$

FIGURE 4. Rates for (a)-(c) mean absorption time and (b)-(d) mean fixation time for all graphs of order 6 under Bernoulli and binomial proliferation. Colour codes are indicated in Figure 3.

Although these graphical experiments are far from describing the whole of dynamics of proliferating processes, they have helped us to perceive the subtlety of temporal monotonicity in $r$ and $p$ and, ultimately, to find a second transition from a regime where proliferation is disadvantageous in terms of fixation time to another where it is advantageous.

### S5. Absorption and fixation curves

Figure 5 allows us to visualize the effect that the increase in population size has on the profiles of mean absorption and fixation times in cliques, cycles and stars under Bernoulli proliferation updating.

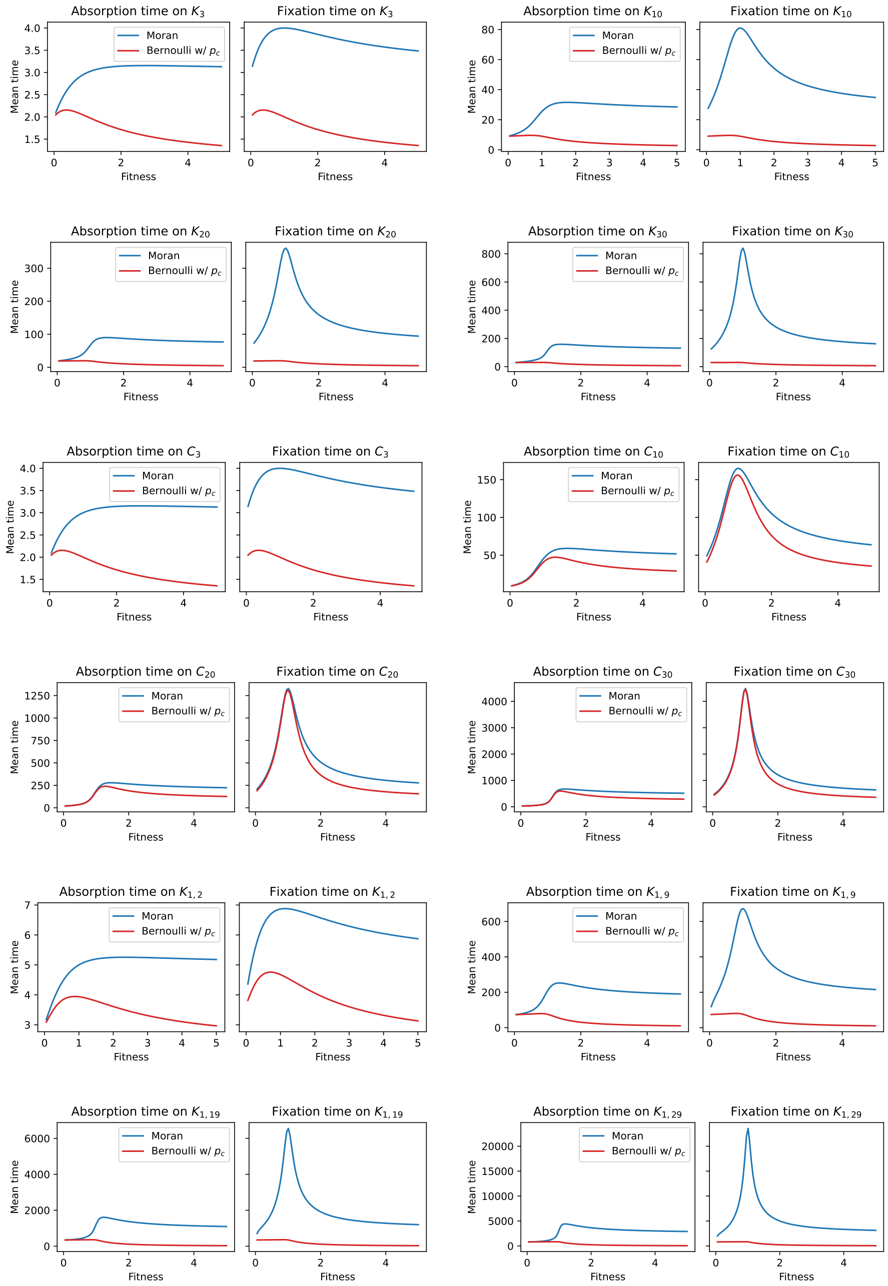


Figure 5. Mean absorption and fixation times on cliques, cycles and stars graphs for Moran and Bernoulli processes when $0 < r \leq 5$ and $p = p_c(r)$.

Universidad de Santiago de Compostela, E-15782 Santiago de Compostela, Spain.

Instituto de Matemática y Estadística Rafael Laguardia, Facultad de Ingeniería, Universidad de la República, J.Herrera y Reissig 565, C.P.11300 Montevideo, Uruguay.

Departamento de Matemáticas, Instituto Universitario de Matemáticas y Aplicaciones (IUMA), Universidad de Zaragoza, Pedro Cerbuna 12, E-50009 Zaragoza, Spain.
